# Historical contingency influences the diversity of feather nanostructures in cuckoos

**DOI:** 10.1101/2023.12.05.570151

**Authors:** Klara K. Nordén, Christopher R. Cooney, Frane Babarović, Mary Caswell Stoddard

**Author notes:** E-mail addresses,.

## Abstract

Structural coloration is widespread in animals, yet we know relatively little about its evolution and development. While previous studies have explored adaptive functions of structural color, a key gap is our lack of understanding of how historical contingency (path-dependency of biological processes) influences the loss and gain of this complex trait. We shed light on this question by describing feather nanostructures responsible for plumage colors in the cuckoos (family Cuculidae), a group with widespread occurrence of shiny, metallic plumage (metallic luster). The melanosomes found in feathers with metallic luster have specialized shapes: hollow rods, thin solid rods, hollow platelets, or solid platelets. In contrast, it is generally assumed that drably colored feathers possess thick, rod-shaped melanosomes. However, we uncover that this assumption is unfounded in cuckoos. We describe metallic luster in the plumages of 126 cuckoo species and map its phylogenetic distribution. This reveals that metallic luster is widespread in cuckoos but has likely been lost several times. We then use transmission electron microscopy to describe the feather nanostructures of 21 cuckoo species. Surprisingly, the drab feathers of many cuckoo species contain melanosomes with specialized shapes. We propose that historical contingency greatly influences nanostructure diversity in cuckoos. Specialized melanosome shapes can be retained in the plumages of drab species, potentially making it easier for metallic luster to evolve again in the future. This discovery supports the idea that historical contingency plays a key role in shaping the evolution of plumage color diversity.

## 1. Introduction

The naturalist Alfred Russell Wallace noted in his landmark paper “The Colors of Animals and Plants” (Wallace, 1877) that clades of animals often share a similar color palette: “We […] find that color is constant in whole genera and other groups of species. The Genistas are all yellow, the Erythrinas all red; many genera of Carabidae are entirely black; whole families of birds—as the Dendrocolaptidae—are brown; while among butterflies the numerous species of Lycaena [Lycaenidae] are all more or less blue, those of Pontia white, and those of Callidryas yellow.” (Wallace, 1877). Birds, which display a vast variety of plumage colors (Stoddard & Prum, 2011), have been a favorite group in which to study color in nature. Following Wallace and other foundational work (e.g. Endler, 1992), the focus has chiefly been on exploring the adaptive functions of plumage colors. For example, plumage color has been shown to evolve in response to sexual selection (Cooney et al., 2019, Dale et al., 2015), environmental lighting conditions (Gomez & Théry, 2004, Marcondes & Brumfield, 2019), background color (crypsis, Troscianko et al., 2016, Mason et al., 2023) and climate (thermoregulation, Romano et al., 2019, Rogalla et al., 2021). A large body of research shows that natural and sexual selection are key forces in shaping animal colors (Cuthill et al., 2017, Hill & McGraw, 2006a).

However, there is a second (not mutually exclusive) general explanation for color variation in animals which has been much less explored—historical contingency. By historical contingency we mean the path-dependency of biological processes (Desjardins, 2011, Blount et al., 2018), i.e. that evolutionary outcomes are biased by the developmental and genetic background of a lineage (Nordén & Price, 2018, Stryjewski & Sorenson, 2017, Price et al., 2000). Historical contingency is particularly relevant for explaining large-scale patterns in plumage color diversity, such as the reoccurring color themes in families noted by Wallace (1877). This is because birds use many different mechanisms to produce color and each mechanism is associated with a particular genetic and developmental framework that takes time to evolve (Hill & McGraw, 2006b, Price-Waldman & Stoddard, 2021). Once a color mechanism has evolved in a clade, it is likely that it will repeatedly be used in response to selection, as opposed to individual species evolving entirely new mechanisms (Nordén & Price, 2018). Conserved genetic and developmental frameworks have been implicated in the repeated evolution of red carotenoid plumage pigmentation in some clades (Thomas et al., 2014, Nordén & Price, 2018, Twyman et al., 2018, Price et al., 2007, Prager & Andersson, 2010).

Here, we investigate whether historical contingency could explain the widespread occurrence of structural barbule colors in Cuculidae by exploring an aspect of the feather nanostructure—melanosome type. Structural colors arise from the interaction of light with a nanostructure (in contrast to pigmentary colors like carotenoids that produce colors from selective absorbance). Structural coloration in birds can be divided into two main types: 1) structural barbule coloration (Figure 1J), which is produced by arrays of melanin-filled organelles (melanosomes) in the barbules, and 2) structural barb coloration (Figure 1K), which is produced by nanostructured “spongy” keratin in the barbs (Prum, 2006). Though the former are often called “iridescent structural colors” and the latter “non-iridescent structural colors”, both types of structural colors do in fact display iridescence (a change in hue with viewing or observation angle, see Nordén et al., 2023, Skigin et al., 2019, Stavenga et al., 2011). However, only structural barbule coloration produce colors with metallic luster—the metallic sheen, for example, of the peacock’s plumage. We have previously shown that metallic luster is characterized by colored specular reflectance and a low diffuse reflectance, which is present in structural barbule but not structural barb coloration (Nordén et al., 2023). Therefore, we will use metallic luster, and not iridescence, to describe structural barbule colors here.

**Figure 1.**
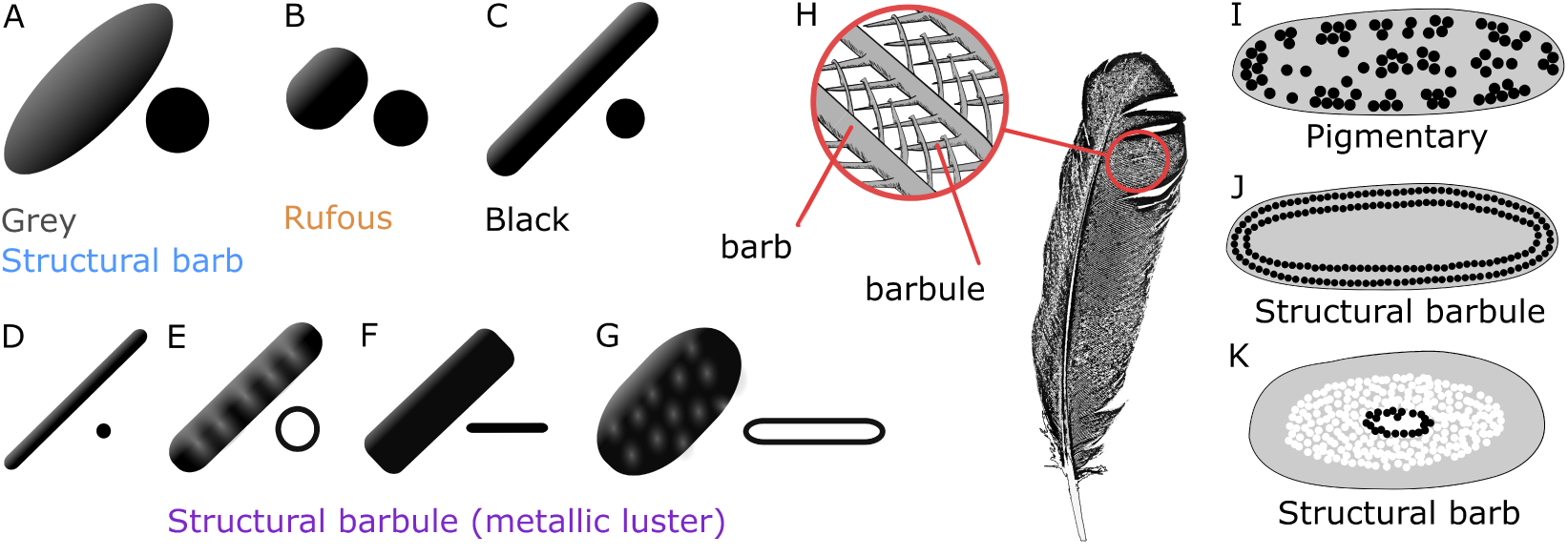
Overview of melanosome types and their structural arrangement in feather filaments. Three main types of melanosomes have been described from pigmentary feathers: (A) large, ellipsoids in gray feathers, (B) short rods in rufous feathers, and (C) thick solid rods (*≥* 190nm in diameter) in black feathers. Thick solid rods can also produce faint (but not intense) metallic luster in some species (Nordén et al., 2021). Four derived melanosome types have been described from feathers with metallic luster: (D) thin solid rods (*<* 190nm in diameter), (E) hollow rods, (F) platelets, and (G) hollow platelets. Melanosomes are also organized differently in the feather (H) depending on color mechanism. Cross-sections of (I) a barbule from a pigmentary feather with disorganized thick solid rods, (J) a barbule from a feather with metallic luster with thin solid rods organized in multiple layers, (K) a barb with nanostructured “spongy” keratin and melanosomes in the interior. Figure adapted from Figure 2 in Nordén et al., 2021, and Figure 2 in Nordén et al. (2023). Note that colors produced from structural barbules are typically called “iridescent structural colors”—here we use metallic luster following the development of this concept by Nordén et al. (2023).

The frequency of metallic luster varies greatly in different bird families, as was noted by the Austrian ornithologist Ludwig Auber over fifty years ago (Auber, 1957). In some families, like Anatidae, Phasianidae, Cuculidae and Columbidae, metallic luster is very frequent and often bright—but in others it is absent from most species, like Psittacidae and Cotingidae. Species in the Psittacidae and Cotingidae families instead use other color mechanisms to produce bright colors (pigments and structural barb coloration (Prum et al., 2012, Tinbergen et al., 2013)). The evolution of bright and saturated metallic luster depends on the evolution of derived melanosome types (thin solid rods, hollow rods, solid platelets or hollow platelets) that differ from the ancestral thick solid rods found in pigmentary black or gray plumage (Figure 1A–G, see Nordén et al., 2021). During feather development, these melanosomes must be arranged into precisely spaced layers within the feather barbule to produce interference colors (Durrer, 1977, Maia et al., 2012). Moreover, feathers with metallic luster often display modified barbules, which are flattened and twisted to allow maximal reflection with the incoming light (Durrer, 1977, D’Alba et al., 2021, Yoshioka & Kinoshita, 2002). Thus, producing bright and saturated metallic luster requires several modifications to melanosome and feather development, and traits depending on the combined effect of multiple modifications are likely more difficult to evolve. We therefore hypothesize that historical contingency is an important factor in explaining the large-scale presence and absence of metallic luster in bird clades.

Intriguing evidence from hummingbirds (family Trochilidae) and tree swifts (family Hemiprocnidae) provides a possible mechanism for the evolution of metallic luster as a historically contingent process. Until recently, derived melanosomes (thin solid rods, hollow rods, solid platelets or hollow platelets, Figure 1D–G) were thought to be uniquely found in plumage with metallic luster. Melanosomes are also present in black, gray, and brown/rufous feathers—but in this case they do not form nanostructures and therefore only function as a pigment (Figure 1I). Melanosomes are typically shaped as thick solid rods in black and gray plumage (Figure 1A, C), and as short rods in rufous plumage (Figure 1B)(Li et al., 2010). Thus, it has been assumed that melanosome type is tied to plumage color production (Nordén et al., 2021, Li et al., 2012, 2010, Vinther, 2020, Nordén et al., 2019). This relationship between melanosome type and color has been used to reconstruct colors of extinct animals, whose integumentary melanosomes sometimes fossilize (Nordén et al., 2019, Li et al., 2010, 2012, Babarović et al., 2019, Hu et al., 2018, Vinther et al., 2008, 2010). However, Smithwick (2019) described hollow platelets extracted from the gray and black plumage of several hummingbird species, and solid platelets from the gray plumage of tree swifts. These families are known to produce hollow platelets and solid platelets respectively in plumage with metallic luster (Durrer, 1977). Hummingbirds and tree swifts are unusual in that not a single species completely lack patches with metallic luster. This led Smithwick (2019) to suggest that derived melanosomes may be retained in feather patches across the entire body plumage if a species has metallic luster in at least one patch. Here, we take Smithwick’s hypothesis one step further and ask whether derived melanosomes may be retained in species that have lost metallic luster altogether. We hypothesize that while there is selective pressure to gain derived melanosomes when evolving metallic luster, there is no selective pressure to lose derived melanosomes when transitioning back to pigmentary colors. This is because the derived melanosomes would function much like a thick solid rod—simply as an absorbing pigment—if the nanostructural order in the barbule was lost. The shape of the melanosome is critical for the production of metallic luster (a structural color), but it is irrelevant for the absorbance properties of the pigment melanin. We speculate that the retention of derived melanosomes could provide a mechanistic explanation for the repeated gains and losses of metallic luster seen in some bird families, which gives rise to the biased distribution of this structural color in bird families noted by Auber (1957). In other words, once derived melanosomes evolve in a clade in order to achieve bright metallic luster in the plumage, this is likely to cause a higher frequency of metallic luster in this clade more generally. Though previous studies have explored the evolution of feather nanostructures in several clades with metallic luster (including starlings (family Sturnidae) (Maia et al., 2013, 2016, Durrer & Villiger, 1970), ducks (family Anatidae) (Eliason et al., 2015), hummingbirds (family Trochilidae) (Gruson et al., 2019), and trogons (family Trogonidae) (Quintero & Espinosa de los Monteros, 2011, Durrer & Villiger, 1966)), no study has yet explored the drably colored species in the same clade. This represents an important missing puzzle piece in our understanding of the loss and gain of this structural color.

We address this gap by exploring the diversity of feather nanostructures in the cuckoos (family Cuculidae), a family with widespread occurrence of metallic luster in the plumage. Previous studies have described three types of derived melanosomes in this clade from species with metallic luster: thin solid rods (in *Chrysococcyx cupreus* (African emerald cuckoo), described by Durrer, 1977), hollow rods (in *Centropus sinensis* (greater coucal), *Centropus violaceus* (violaceous coucal) and *Centropus ateralbus* (white-necked coucal), described by Nordén et al., 2019) and solid platelets (in *Phaenicophaeus diardi* (black-bellied malkoha) and *Phaenicophaeus curvirostris* (chestnut-breasted malkoha), described by Nordén et al., 2019). We first phylogenetically mapped the presence of metallic luster in 126 cuckoo species (excluding 21 species lacking genetic data for the phylogenetic reconstruction) using a three-point scale based on visual assessment of their plumage. We then selected 21 representative species across the tree with and without metallic luster, and measured their plumage colors with cross-polarization photography to validate our visual scoring and quantify differences in metallic luster. From these representative species, we imaged crosssections of feather samples using transmission electron microscopy (TEM) to compare melanosome type and feather nanostructure in species with and without metallic luster in five main clades. Synthesizing these approaches with data on melanosome shape diversity which has previously been described across all birds, we described the evolution and diversity of feather nanostructures in Cuculidae and explored the relationship between melanosome type and plumage color.

## 2. Results

### 2.1. Evolution of metallic luster in Cuculidae

We assigned scores to 10 plumage patches (crown, nape, mantle, rump, dorsal tail, wing covert, wing primaries/secondaries, throat, breast, and belly) to quantify the presence of metallic luster in cuckoos. Since metallic luster can vary in intensity (brightness and saturation), we scored it on a three-point scale: 0: absence, 1: faint-moderate, 2: intense (see §4.1, *Methods*, for more details). We validated our scores on a subset of samples using cross-polarization photography, a technique that can measure the strength and saturation of the specular reflection, which we use as a proxy for metallic luster (see Nordén et al., 2023). A high relative specular reflection that has some saturation (is colored) indicates presence of metallic luster, while low relative specular reflection and/or an unsaturated specular reflection indicates absence of metallic luster. The sampled patches did indeed vary in metallic luster, with patches scored as “0” measuring low relative specular reflectance and saturation and patches scored as “1” or “2” with higher specular reflection and saturation (Figure 2), when measured with cross-polarization photography. While there is some overlap in “1” and “2”, in general “2” has the highest specular reflection and specular saturation (Figure 2).

**Figure 2.**
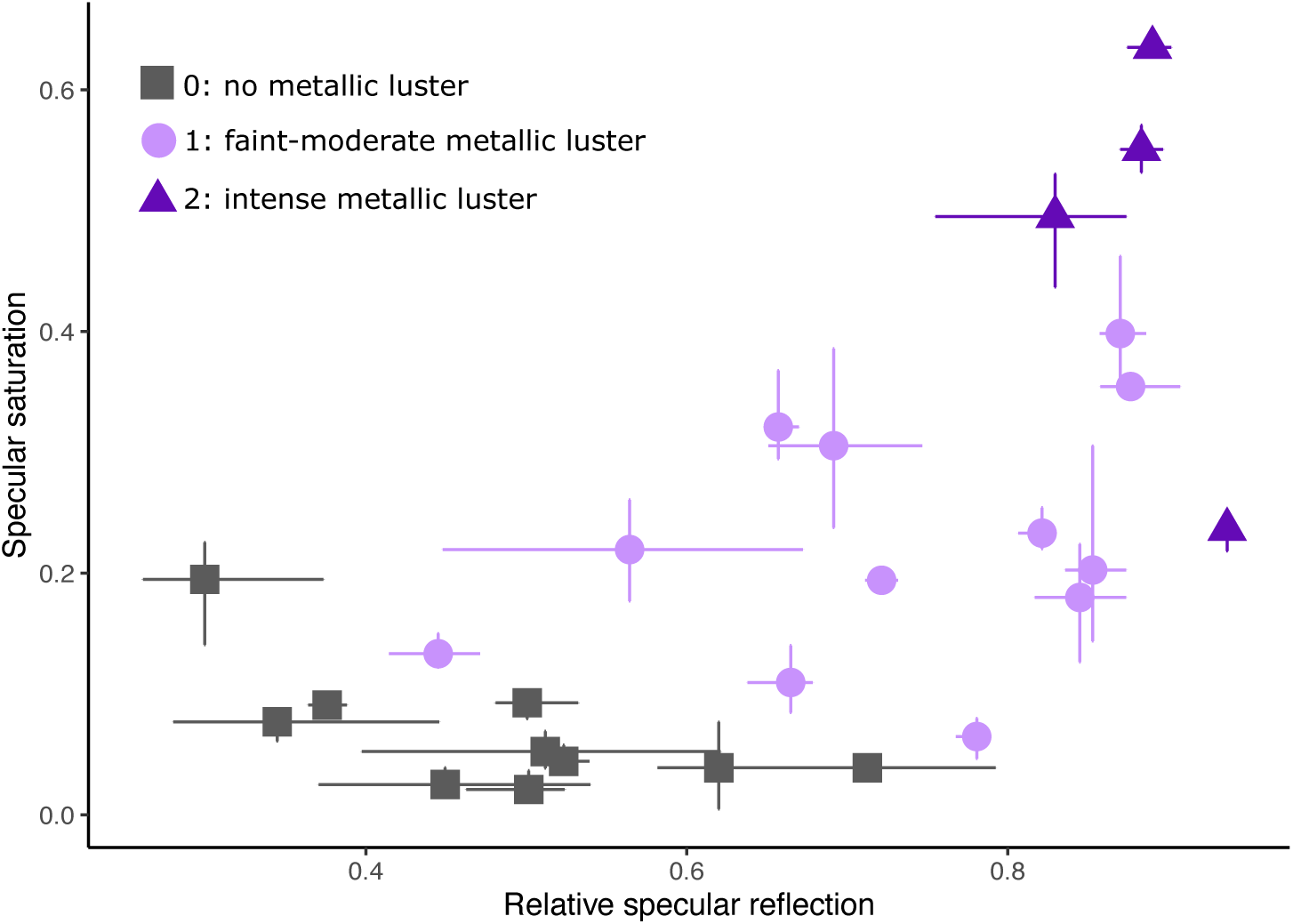
Plumage patches sampled from Cuculidae species vary in the intensity of metallic luster, as measured by the relative specular reflection and specular saturation. These measures were extracted using cross-polarization photography (see Methods for details). Colors and shapes indicate the score for the plumage patch based on visual assessment. Patches scored as “0” tended to have low relative specular reflection and saturation, indicative of absence of metallic luster, while patches scored as “1” and “2” recorded higher relative specular reflection and saturation, indicative of faint-intense metallic luster. “1” and “2” show overlapping distributions, but in general “2” measures the highest relative specular reflection and saturation. Lines represent the range of three repeated measurements of the same patch.

Metallic luster is widespread in Cuculidae, with over three-fifths of species (62%) exhibiting the trait in at least one plumage patch (Figure 3A). Faint-moderate metallic luster is most common, while intense metallic luster is limited to a single genus (*Chrysococcyx*, Figure 3A, E). In terms of distribution on the body, metallic luster is more frequent in the dorsal than the ventral part of the plumage (Figure 4). Since the nanostructures giving rise to metallic luster are built with melanin, this pattern may be linked to the general bias for a melanized dark ventral and light dorsal body seen in many animals (for which the evolutionary reason is still debated, Rowland, 2008). Alternatively, or additionally, ventral plumage may tend to lack metallic luster because flattened barbules—which is a common barbule modification in plumage with metallic luster (Durrer, 1977)—may decrease waterproofing of feathers (Eliason & Shawkey, 2011).

**Figure 3.**
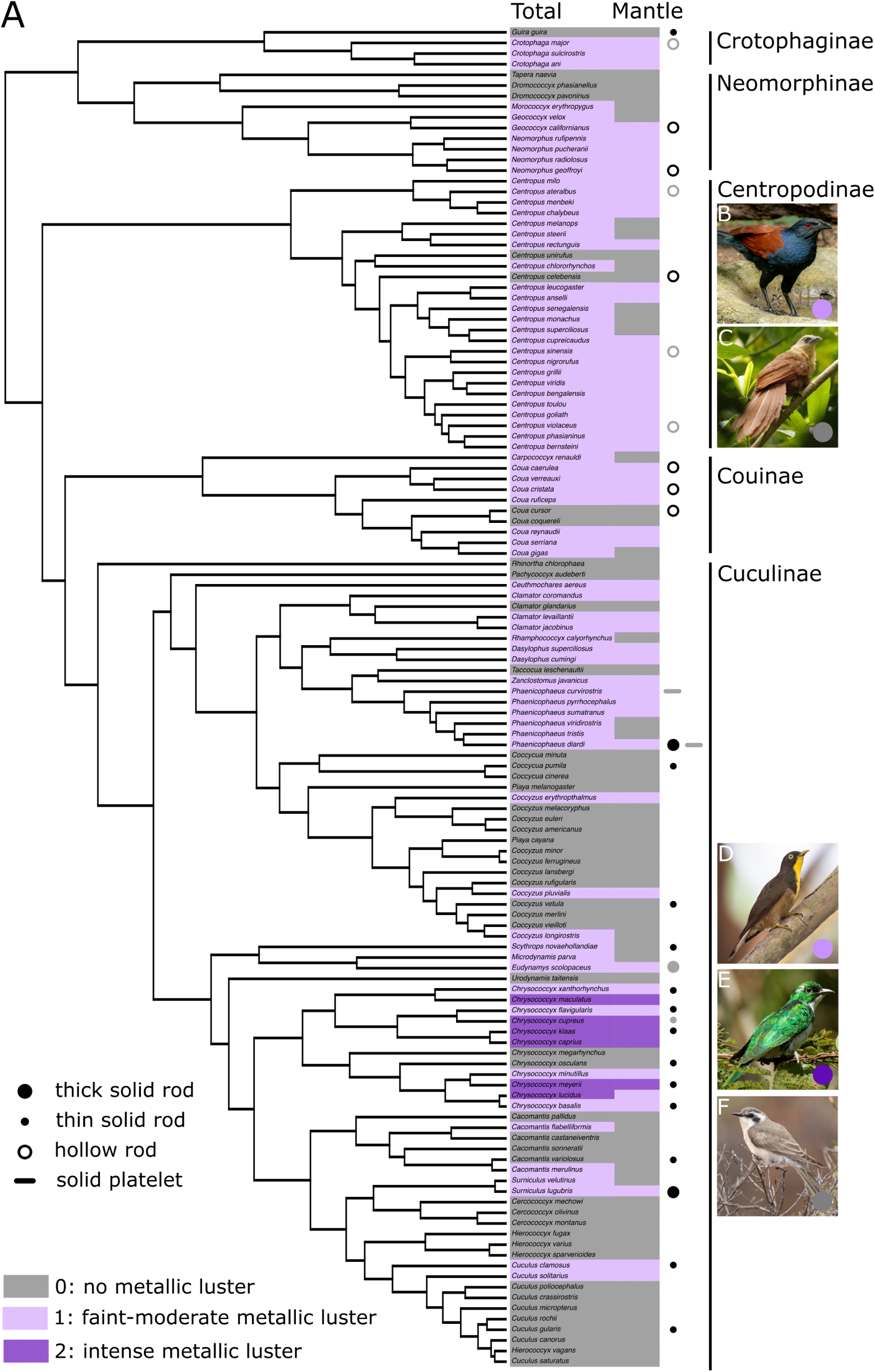
(Previous page) Plumage with metallic luster is widespread in Cuculidae. Each species was scored for the presence of metallic luster on a three-point scale (0: absent; 1: faint-moderate; 2: intense) across body patches (see §4.1, *Methods*). The result of this analysis is color coded on the tree (A) with gray, light and dark purple representing 0, 1 and 2 scores, respectively. “Total” describes the highest score received by each species anywhere on the body, “mantle” describes the score for the mantle patch. Melanosome type in the plumage is represented by symbols (large dot, thick solid rods; small dot, thin solid rods; open circle, hollow rod; platelet, solid platelet)—black symbols represent melanosome types described from feather samples in this study, and gray symbols represent types described in previous studies (*Chrysococcyx cupreus* described in Durrer & Villiger, 1970, all others from Nordén et al., 2019). The photographs showcase the variation in metallic luster of the cuckoos we sampled: B) *Centropus sinensis* (all rights reserved by copyright holder Sakkarin Sansuk, reproduced here with permission) C) *Centropus celebensis* (all rights reserved by copyright holder Marc Thibault, reproduced here with permission), D) *Chrysococcyx flavigularis* (all rights reserved by copyright holder Dubi Shapiro, reproduced here with permission), E) *Chrysococcyx klaas* (all rights reserved by copyright holder Luke Seitz, reproduced here with permission), F) *Chrysococcyx osculans* (all rights reserved by copyright holder David Ongley, reproduced here with permission).

**Figure 4.**
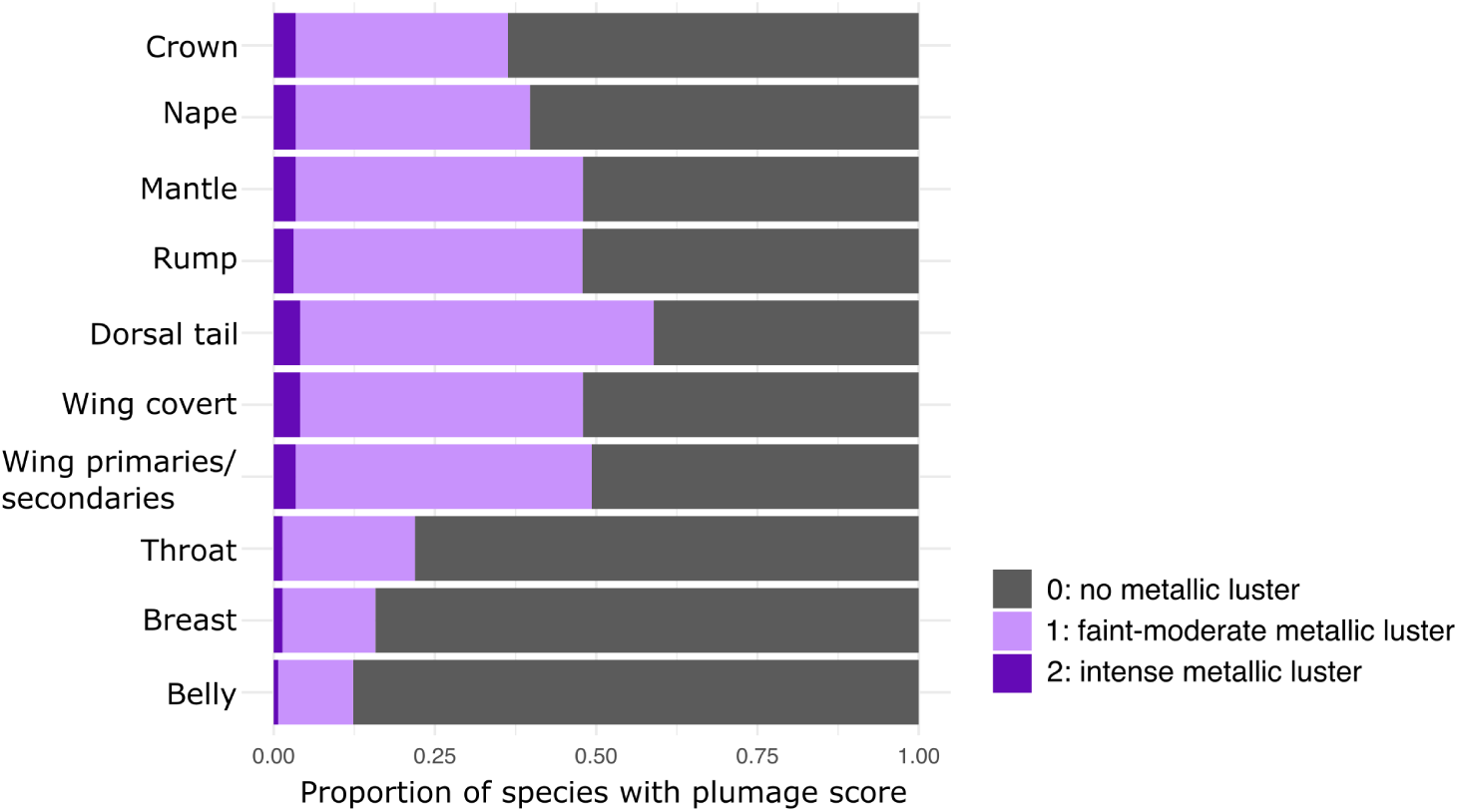
Metallic luster is more common in the dorsal (crown, nape, mantle, rump, dorsal tail, wing coverts, and wing primaries/secondaries) compared to the ventral plumage (throat, breast, and belly) in the Cuculidae. Bars show the proportion of species scored as “0”, “1” or “2” (gray, light purple and dark purple respectively) for each plumage patch.

Nine genera lack metallic luster completely (*Guira*, *Tapera*, *Dromococcyx*, *Rhinortha*, *Pachycoccyx*, *Coccycua*, *Urodynamus*, *Cerococcyx* and *Hierococcyx*), but most genera contain species exhibiting a mix of patches with and without metallic luster. In some cases, species lacking metallic luster appear nested within a clade exhibiting the trait (in one or more patches)—e.g. *Coua cursor* (running coua) and *Coua coquereli* (Coquerel’s coua) within the *Coua* clade, and *Centropus celebensis* (bay coucal)(Figure 3C) and *Centropus unirufus* (rufous coucal) within the *Centropus* clade. Similarly, *Chrysococcyx flavigularis* (yellow-throated cuckoo)(Figure 3D) exhibits faint-moderate metallic luster but is nested in a clade with intense metallic luster (Figure 3A, E). In other cases,the situation is reverse: species with metallic luster appear nested within clades otherwise lacking the trait—e.g. *Coccyzus pluvialis* (chestnut-bellied cuckoo) within the *Coccyzus* clade, and *Cacomantis flabelliformis* (fan-tailed cuckoo) within the *Cacomantis* clade (Figure 3A). We cannot make definitive inferences from these patterns, but based on phylogenetic bracketing, these are potential cases of losses and gains of metallic luster, respectively. While we did attempt to model rates of character transitions and reconstruct ancestral states using Bayesian inference and continuous-time Markov models, our results were highly variable. This suggests that gains and losses of metallic luster are not well represented by a Markov process and/or are difficult to reconstruct with existing approaches, and we therefore chose to not include the results here (see Appendix A for further discussion).

Based on this overview of the distribution of metallic luster in Cuculidae, we can start to explore the nanostructural diversity within feathers with and without metallic luster. If derived melanosomes are retained when metallic luster is lost, we should observe this in species lacking metallic luster nested within clades that have metallic luster. We turn to this topic in the next section (§2.2, *Nanostructural diversity*).

### 2.2. Nanostructural diversity

To characterize feather nanostructural diversity, feather samples were collected from museum specimens representing 21 species across the Cuculidae. All feather samples were obtained from the mantle patch to facilitate direct comparisons between species (with the exception of *Phaenicophaeus diardi*, which was sampled from the belly patch, and *Chrysococcyx xanthorhynchus* (violet cuckoo) from which a loose feather of unknown patch was opportunistically sampled). Samples were embedded in resin blocks, sectioned and imaged under transmission electron microscopy to explore feather nanostructure. All patches sampled were part of the set photographed with cross-polarization to validate differences in metallic luster (Figure 2). We also imaged all feather samples with confocal microscope to document the macro-scale appearance of the feather.

#### 2.2.1. Plumage with metallic luster

We first describe the melanosome type and organization found in species with metallic luster (scored as “1” or “2”).

##### Melanosome type

We found that plumage with metallic luster is produced with either solid rods (in *Chrysococcyx*, *Surniculus* and *Cuculus*, Figure 5A–G, Figure 3A) or hollow rods (in *Neomorphus*, *Geococcyx* and *Coua*, Figure 6A–D, Figure 3A). Using image analysis to identify melanosome cross-sections in microscope images (see §4.3 for detailed methods), we measured the diameter of solid rods in imaged barbule cross-sections to determine whether they could be considered thick solid rods (*≥* 190nm in diameter) or derived thin solid rods (*<* 190nm in diameter, see Nordén et al., 2021). We found that melanosomes in all species sampled could be considered to be thin solid rods, except in *Surniculus lugubris* (square-tailed drongo-cuckoo), which had melanosomes with an average diameter of 203nm (Figure 7A, Figure 5F). From a previous study (Nordén et al., 2019), we also know that hollow rods are present in the plumage patches with metallic luster in *Crotophaga major* (greater ani), *Centropus sinensis*, *Centropus violaceus* and *Centropus ateralbus*, while solid platelets are present in *Phaenicophaeus diardi* (Figure 5H) and *Phaenicophaeus curvirostris*. Since melanosome type in plumage with metallic luster is often conserved within a genus (Nordén et al., 2021, Durrer, 1977, Quintero & Espinosa de los Monteros, 2011), these observations suggest that metallic luster is produced with hollow rods in the clades Crotophaginae, Neomorphinae, Centropodinae and Couinae, while Cuculinae produces metallic luster with thin solid rods (with the exception of *Phaenicophaeus*). A summary of the known melanosome types in the cuckoos is shown in Figure 3A.

**Figure 5.**
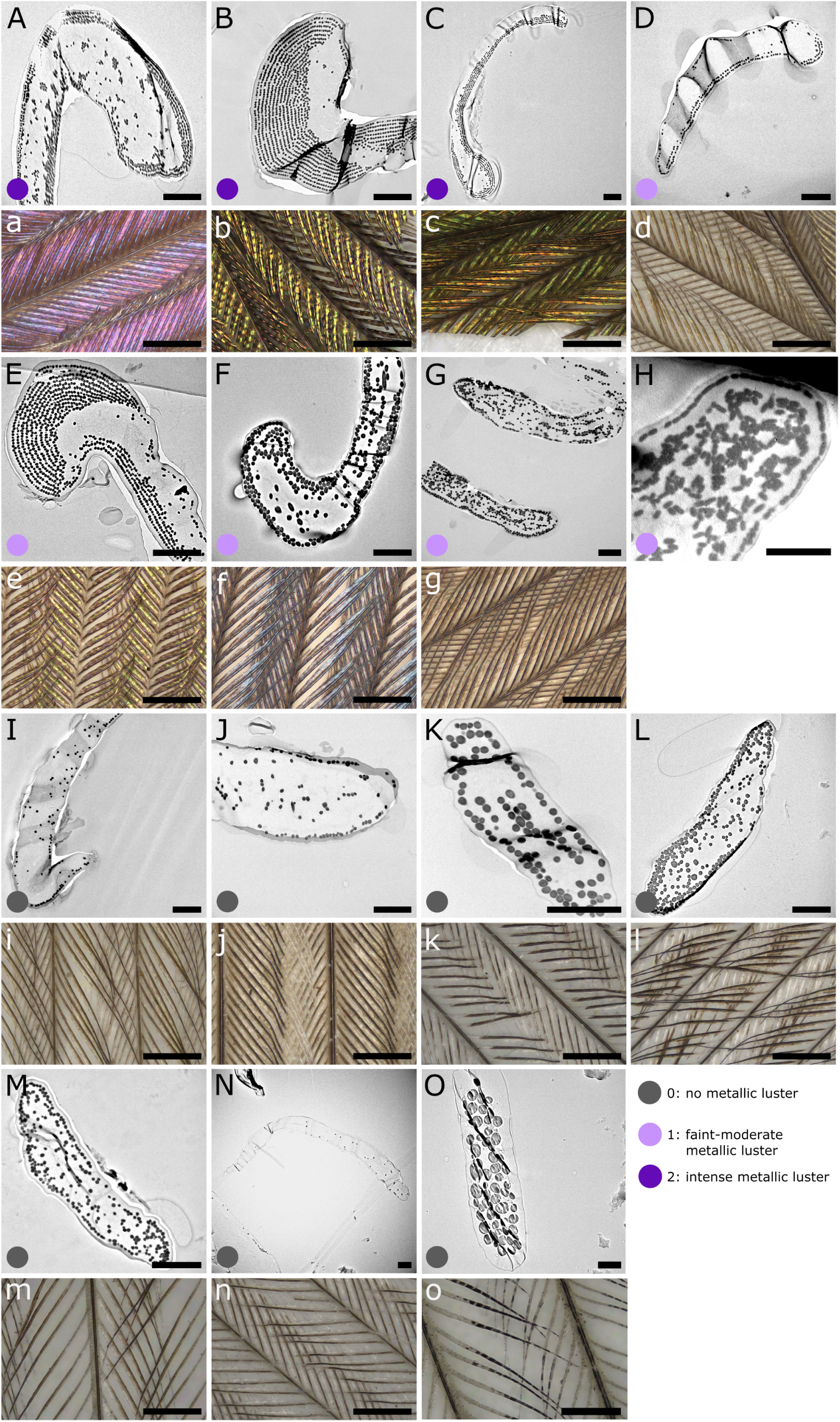
(Previous page) Feather nanostructures in plumage with and without metallic luster in the Cuculinae clade. Dark and light purple dots indicate intense and faint-moderate metallic luster, respectively. Gray dots indicate absence of metallic luster. Capital letters denote transmission electron microscope images of barbule cross-sections; lower case letters denote confocal microscope images of each feather sample. A-a) *Chrysococcyx xanthorhynchus*, B-b) *Chrysococcyx klaas*, C-c) *Chrysococcyx meyeri*, D-d) *Chrysococcyx basalis*, E-e) *Chrysococcyx flavigularis*, F-f) *Surniculus lugubris*, G-g) *Cuculus clamosus*, H) *Phaenicophaeus diardi* (from Nordén et al., 2019), I-i) *Chrysococcyx osculans*, J-j) *Scythrops novaehollandiae*, K-k) *Cuculus gularis*, L-l) *Cacomantis variolosus*, M-m) *Coccyzus vetula*, N-n) *Coccycua pumila*, O-o) *Phaenicophaeus diardi*. All scale bars in TEM (A-G) and confocal (a-g) images equal 2*µ*m and 200*µ*m, respectively.

**Figure 6.**
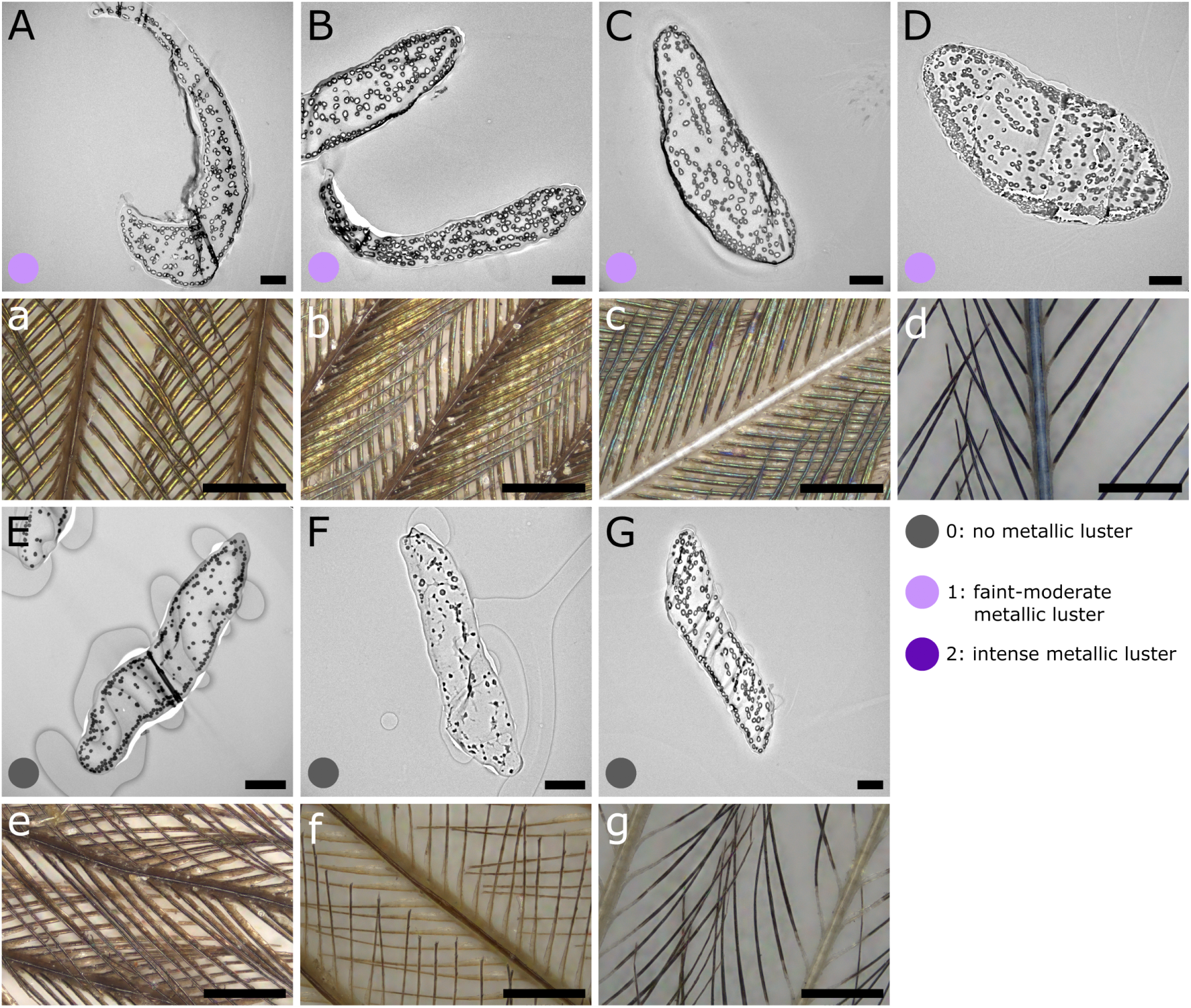
Feather nanostructures in plumage with and without metallic luster in the Crotophaginae, Neomorphinae, Centropodinae and Couinae clades. Dark and light purple dots indicate intense and faint-moderate metallic luster, respectively. Gray dots indicate absence of metallic luster. Capital letters denote transmission electron microscope images of barbule cross-sections; lower case letters denote confocal microscope images of each feather sample. A-a) *Neomorphus geoffroyi*, B-b) *Geococcyx californianus*, C-c) *Coua cristata*, D-d) *Coua caerualela*, E-e) *Guira guira*, F-f) *Centropus celebensis*, G-g) *Coua cursor*. All scale bars in TEM (A-G) and confocal (a-g) images equal 2*µ*m and 200*µ*m, respectively.

**Figure 7.**
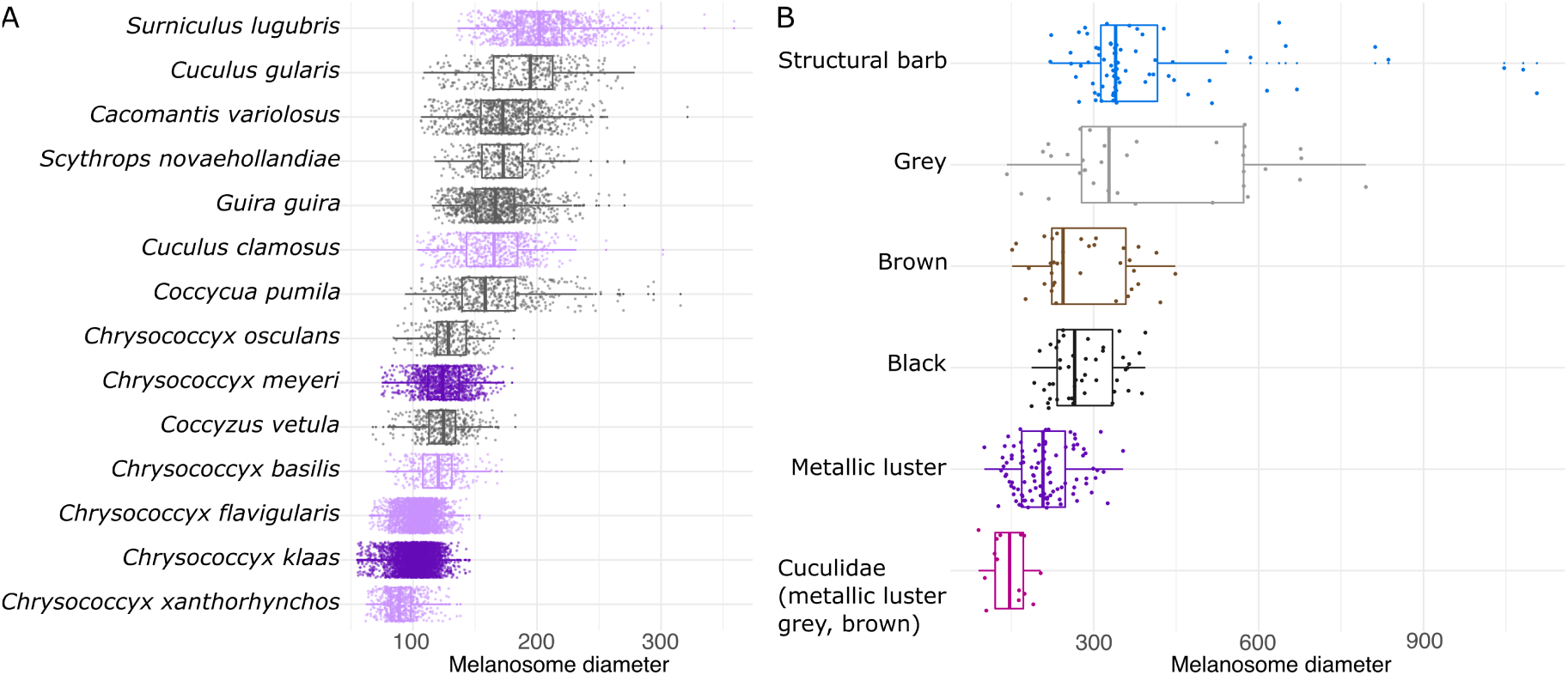
The diameter of solid rods in the Cuculidae is small irrespective of plumage color (A), which contrasts with the diversity seen across plumage colors in a broader sample of birds (B). Data in B (except for Cuculidae) from the following studies: Li et al. (2012), Babarović et al. (2019), Nordén et al. (2019).

##### Melanosome organization

We found that in all feathers with metallic luster, melanosomes form a single or multiple ordered layers towards the outer border of the barbule (Figure 5A–H; Figure 6A–D). Most structures have only a single outer layer, while species in the genus *Chrysococcyx* exhibited up to 10 layers of melanosomes (Figure 5A–E). A greater number of layers increases the total reflection of the structure, resulting in brighter and more saturated colors (Kinoshita et al., 2008). Interestingly, *Chrysococcyx flavigularis* (Figure 5E), a species with faint-moderate metallic luster in the mantle patch, has feathers containing as many melanosome layers as its sister species *Chrysococcyx klaas* (Klaas’s cuckoo) (Figure 5B), a species with intense metallic luster in the mantle patch. However, closer inspection reveals that the melanosome layers in *Chrysococcyx flavigularis* are uneven and show a higher degree of disorder than in *Chrysococcyx klaas*. This demonstrates that even a slight decrease in nanostructural order—while retaining melanosome type—can cause a stark change in plumage color.

#### 2.2.2. Plumage without metallic luster

Having surveyed melanosome diversity in cuckoo species with metallic luster, we can now contrast this with the melanosome type and organization in species lacking metallic luster.

##### Melanosome type

Derived melanosome types (deviations from the thick solid rods that are ancestral in birds) were found in all but one of the plumage samples lacking metallic luster that we studied (the exception being the gray belly of *Phaenicophaeus diardi*). Moreover, the species lacking metallic luster had retained the same derived melanosome type that were present in species with metallic luster of that clade. In *Coua* and *Centropus*—two genera where species with metallic plumage have hollow rods—hollow rods were also present in the gray plumage of *Coua cursor* (Figure 6G) and the rufous plumage of *Centropus celebensis* (Figure 6F). In *Centroupus celebensis*, hollow rods are interspersed with irregularly shaped melanosomes, which may indicate a disruption of melanosome formation (see Li et al., 2019)(Figure 6F). In *Chrysococcyx* —a genus where species with metallic plumage have thin solid rods—thin solid rods were also found in the gray plumage of *Chrysococcyx osculans* (black-eared cuckoo). Even species in genera that mostly lack metallic luster were found to have thin solid rods in their plumage: *Coccycua pumila* (dwarf cuckoo), *Coccyzus vetula* (Jamaican lizard cuckoo), *Cuculus gularis* (African cuckoo), *Cacomantis variolosus* (brush cuckoo), *Scythrops novaehollandiae* (channel-billed cuckoo) (Figure 5J–N) and *Guira guira* (guira cuckoo) (Figure 6E). The exception was the gray belly feathers of *Phaenicophaeus diardi*, which had large melanosomes (approximately 1*µ*m in diameter) that were likely ellipsoid in shape, based on the varied diameter seen in the cross-sections (Figure 5O). Thus, they were markedly different from the solid platelet melanosomes previously described in the mantle of *Phaenicophaeus diardi* with metallic luster (Figure 5H)(Nordén et al., 2019), and more similar to the melanosomes that have previously been described in gray plumage of other birds (Figure 1A).

##### Melanosome organization

Melanosomes in samples without metallic luster tended to form a layer at the outer border of the barbule (Figure 5I–N, Figure 6E, G), similar to the melanosome organization seen in samples with metallic luster. However, this layer had multiple gaps (Figure 5I–N, Figure 6E, G). In general, we observed that the number of melanosomes in barbules from feathers without metallic luster appeared to be lower than those in feathers with metallic luster. This was sometimes seen directly in individual cross-sections (e.g. Figure 5I, J and N), and in other cases seen from the distribution of melanin in the feather as a whole. Melanin was not deposited in the full length of the barbule but rather only in a section of the barbule (Figure 5i–n, Figure 6f–g). Similar patterns of partial melanization have been described in the barbs of gray down (Dove, 2000). Though difference in melanin content would have to be confirmed with a chemical test, we note that the plumage we sampled without metallic luster was gray in color, while plumage with metallic luster is much darker (in transmitted light). Thus, we speculate that the decreased number of melanosomes may be a key factor disrupting the production of metallic luster, since not enough melanosomes are available to form continuous nanostructures in the entire barbule. This is supported by previous studies which have suggested that melanosome concentration is important for nanostructure formation (Maia et al., 2012, D’Alba et al., 2021).

An exception in melanosome organization was seen in the sample from the gray belly of *Phaenicophaeus diardi*, which was also the only sample with large, ellipsoidal melanosomes (Figure 5O). Here, instead of forming a layer at the outer border of the barbule, the melanosomes grouped together at the center of the barbule.

#### 2.2.3. Summary

Derived melanosomes are widespread in Cuculidae irrespective of plumage color (metallic luster versus gray or brown, Figure 3A). In particular, we found three instances in which species lacking metallic luster nested in clades with metallic luster had the same derived melanosome types as the parent clade (*Coua cursor* (Figure 6G), *Centropus celebensis* (Figure 6F), and *Chrysococcyx osculans* (Figure 5I)). This suggests that the derived melanosome types in these groups have been retained even as metallic luster was lost.

Derived melanosomes in plumage without metallic luster still formed partial layers in the outer border of the barbule, in a similar manner to melanosomes in plumage with metallic luster (Figure 5, Figure 6). However, the concentration of melanosomes appeared to have been reduced, resulting in partial melanization of the barbules. Thus, metallic luster may be lost by changing the concentration of melanosomes in the feather barbule, which in turn disrupt the nanostructure formation, rather than reverting melanosome type back to an ancestral thick solid rod. A lower concentration of melanosomes is one way of producing gray plumage color, and could therefore explain why many species lacking plumage with metallic luster in the Cuculidae have gray, as opposed to black, plumage.

*Phaenicophaeus diardi* differed from the pattern described above. It had large, ellipsoidal melanosomes in its gray plumage and solid platelets in the mantle patch with metallic luster (Figure 5H, O). Without also sampling the belly patch of other species, it is hard to say if *Phaenicophaeus diardi* is an exception to other cuckoos in terms of how it produces gray color, or an indication that melanosome type varies with body patch.

#### 2.2.4. Decoupling between melanosome type and plumage color

The melanosome diversity in plumage with and without metallic luster that we have described in the section above (§2.2, *Nanostructural diversity*) suggest a decoupling between melanosome type and plumage color. This is in contrast to previous studies, which have shown a relationship between melanosome type and plumage color based on melanosome shape parameters collected from plumage samples of over 400 species (Li et al., 2010, 2012, Nordén et al., 2019, Babarović et al., 2019). To visualize this decoupling more clearly, we can compare the diameter of solid rods from different plumage colors recorded across birds to that of solid rods in Cuculidae (Figure 7B). It is clear that the variation of melanosome diameter in Cuculidae is considerably smaller than what we would expect given their color variability (including metallic luster and gray color, Figure 7B). The diameter of the solid rods in Cuculidae differs significantly from all other color categories except metallic luster (ANOVA one-way test: *df* = 5, *F* = 33.23, *p <* 0.001; Tukey’s post-hoc test, all comparisons *p <* 0.005 except for metallic luster, *p* = 0.41).

## 3. Discussion

We have described the evolution of metallic luster and its relationship to nanostructural barbule variation in the Cuculidae family. A key finding is that derived melanosome types can be retained in species lacking plumage with metallic luster. This is interesting because it suggests that derived melanosomes, once evolved, can be retained over long evolutionary timescales, potentially providing a mechanism for re-emergence of the trait. The widespread occurrence of derived melanosomes across Cuculidae—even in clades where the majority of species lack metallic luster (e.g. *Cuculus*, *Coccycua* and *Coccyzus*, Figure 7A)—suggests that derived melanosomes evolved early in the clade and have then been retained. Rather than switching melanosome type, species that lost metallic luster likely did so by disrupting the nanostructural organization of melanosomes in the barbules. We suggest that this disruption of nanostructural order is driven by a reduction in the number of melanosomes in the feather barbule (Figure 6, 5). Since a lower melanosome concentration and/or partial melanization of barbules is one way of producing gray plumage colors, this might explain why many of the species in Cuculidae lacking metallic luster have gray, as opposed to black, plumage. This is in line with earlier research which has indicated that melanosome concentration in the barbules is important for nanostructure formation (Maia et al., 2012). Thus, we speculate that metallic luster and derived melanosomes evolved early in the Cuculidae clade and that the retention of derived melanosomes in plumage lacking metallic luster has provided a mechanism for the frequent evolution of metallic luster in Cuculidae.

The finding that derived melanosome types can be retained in plumage lacking metallic luster decouples the direct relationship between plumage color and melanosome type that has been shown previously (Figure 1)(Li et al., 2010, 2012, Vinther, 2020, Nordén et al., 2019, Babarović et al., 2019). However, it also validates some of the ideas behind it; specifically the idea that melanosome shape is important for nanostructure formation (Maia et al., 2012). Thin solid rods and hollow rods were seen to form partial layers at the outer border of the barbule even in plumage lacking metallic luster (in *Chrysococcyx osculans*, *Scythrops novaehollandiae*, *Cuculus gularis*, *Cacomantis variolosus* and *Coccyzus vetula*, Figure 5I–N; as well as *Guira guira* and *Coua cursor*, Figure 6E, G). In contrast, the large, ellipsoidal melanosomes in the gray plumage of *Phaenicophaeus diardi* formed clusters towards the middle of the barbule (Figure 5O)—a phenomenon previously described in gray feathers by Babarović et al. (2019). These observations support earlier suggestions that melanosome shape plays a key role in the nanostructural arrangement of melanosomes in the barbule during feather development (Maia et al., 2012, Babarović et al., 2019). However, because there are other parameters that can be adjusted to mute the production of metallic luster—such as the number of melanosomes deposited— evolving a particular melanosome type is not necessary to produce gray plumage color. The great range of melanosome diameters recorded from gray plumage across birds indicates that gray color indeed is produced by a diversity of melanosome types in different species (Figure 7B). This diversity may signal that other clades which, like Cuculidae, contain many species with metallic luster in the plumage (e.g. Anseriformes), also retain derived melanosome types in gray plumage. Such a general pattern of derived melanosome retention in birds would on the one hand complicate direct inferences of plumage color from melanosome types in fossils, but on the other hand give deeper insights into the plumage color evolution of a clade. If derived melanosome types only evolve to produce metallic luster, the finding of such melanosomes in a species would signal the presence of metallic luster in the lineage, albeit not necessarily predicting metallic luster in a particular species.

Evidence that retention of derived melanosomes in plumage lacking metallic luster is more widespread is found in studies of plumage colors in starlings (family Sturnidae). Starlings, like cuckoos, frequently exhibit metallic luster in the plumage and are known to have evolved derived melanosomes (Durrer & Villiger, 1970, Maia et al., 2013, Craig & Hartley, 1985). Though derived melanosomes have not yet been described from starling species that completely lack metallic luster, they have been described from gray and rufous patches in species that also have patches with metallic luster. In *Lamprotornis superbus* (superb starling), the non-metallic rufous plumage on the belly contains hollow platelets—the same melanosome type also present in the species’ metallic dorsal plumage (Rubenstein et al., 2021). Interestingly, many of the melanosomes in the rufous plumage appear irregularly shaped, just like in the rufous plumage of *Centropus celebensis* described in this paper (Figure 6F). These irregularly shaped melanosomes may signal a different melanosome chemistry (greater amount of pheomelanin) and/or a disruption of melanosome shape formation, as indicated by a similar phenomena in chicken (*Gallus gallus*, see Li et al., 2019). Two other species in the same genus, *Lamprotornis fischeri* (Fischer’s starling) and *Lamprotornis unicolor* (ashy starling), have hollow rods in their metallic plumage (Maia et al., 2013) and also exhibit disorganized hollow rods in the gray body plumage (Craig & Hartley, 1985). Intriguingly, losses of metallic luster seem to coincide with transitions to gray or rufous (as opposed to black) melanin coloration in Sturnidae, much like in Cuculidae. This might indicate shared developmental processes involved in the loss of metallic plumage in the two clades.

Our findings add to the growing empirical evidence that historical contingency is important for plumage color evolution (Nordén & Price, 2018, Stryjewski & Sorenson, 2017, Price et al., 2000). An increasingly documented phenomenon directly supporting this argument is color masking in plumage (Price-Waldman & Stoddard, 2021), where one color mechanism masks the effect of a co-occurring mechanism. This can arise if a broadly absorbing pigment (e.g. melanin) masks the effect of another pigment (e.g. carotenoid) or structural barb coloration (Nero, 1954, Moreau, 1958, Hudon et al., 2015, M. Hofmann et al., 2007, Aguillon et al., 2021, D’Alba et al., 2012, Fan et al., 2019, Driskell et al., 2010, Justyn & Weaver, 2023). Though this phenomenon could have adaptive explanations (for example, if the pigment deposition provides other physiological or functional advantages), historical contingency often seems more likely as an explanation. For example, Driskell et al. (2010) found that some species of fairywrens (family *Maluridae*) had spongy, nanostructured keratin in the barbs of black feathers. Spongy keratin in the barbs (Figure 1K) typically gives rise to blue structural coloration (Prum, 2006)—but in the fairy-wrens, the blue color was masked by melanin deposition (Driskell et al., 2010). Since not all species with black plumage exhibited spongy keratin in the barbs, and structural blue coloration is very common in this family, Driskell et al. (2010) concluded that the spongy keratin in black feathers was likely inherited from an ancestral species with blue structural coloration in this patch. Similarly, the red chest and belly of *Passerina ciris* (painted bunting) is produced by carotenoid pigmentation, yet the same feathers also contain spongy, nanostructured keratin in the barbs (Justyn & Weaver, 2023). In this case, the ancestors of the painted bunting likely had full structural blue body coloration (Martínez–Meyer et al., 2004). The widespread occurrence of color masking in bird plumage suggests that transitions between colors can be achieved by switching on or off the masking color in a patch, rather than evolving the color mechanism anew. Thus, we argue that the widespread phenomenon of color masking suggest that historical contingency is an important force in shaping the diversity of plumage colors.

While our study underlines the importance of historical contingency in shaping color evolution and nanostructural diversity, there are also interesting exceptions to this pattern. In *Phaenicophaeus diardi*, solid platelet melanosomes were not retained in the gray belly patch, despite being present in the mantle patch with metallic luster (Figure 5H, O). Why did this species differ from all other cuckoos we sampled? One possibility is that melanosome type in the belly patch generally differs from that of the mantle patch in cuckoos—perhaps linked to different feather structures found in these two patches. Another possibility is that transitions from solid platelets to thick solid rods is easier than transitions from other types of derived melanosomes (thin solid rods and hollow rods) to thick solid rods. To resolve this question, and to get a deeper understanding of how metallic luster evolves in birds, studies of the genetics regulating melanosome shape and structuring are key. This might reveal whether the genetic underpinnings of metallic luster in *Phaenicophaeus diardi* differs from that of other cuckoos species. The literature on the genetic basis of structural coloration is small but rapidly growing (reviewed in Price-Waldman & Stoddard (2021), Saranathan & Finet (2021)), and offers a promising path forward for future research. The varied feather nanostructures we have described in Cuculidae, which represents a continuum from no metallic luster to intense metallic luster, provide an opportunity to link phenotype to genotype, with attention to candidate genes previously linked to melanosome shape and structuring (Rubenstein et al., 2021, Gao et al., 2018, Hellström et al., 2011, Li et al., 2019).

## 4. Methods

### 4.1. Distribution of metallic luster in Cuculidae

Metallic luster (in previous literature typically called “iridescent structural color”, see Nordén et al. (2023)) in plumage varies from a faint sheen to bright and saturated colors. We scored this continuous variation on a 0-2 scale to uncover large-scale patterns in the evolution of structural colors in Cuculidae. Scoring was based on illustrations and photographs from Birds of the World (Billerman et al., 2022). Since a single photograph may be misinterpreted due to for example unusual lighting conditions or inaccurate white balance, we always ensured metallic luster was visible in at least two separate photographs before assigning a score. We also used the verbal description of the species to support our interpretation. For example, descriptions such as “glossed green” or “metallic” would suggest metallic luster. The scale we used is a coarser version of a similar qualitative scale developed by Durrer (1977) and Auber (1957), where our score 1 spans “schwacher Schiller/faint iridescence” to “mässiger Schiller/moderate iridescence”, and 2 spans “intensiver Schiller/intense iridescence” to “brillianter Schiller/luxuriant iridescence”. A score of 0 means no metallic luster was observed in the plumage. All scores were assigned by K. K. N., based on their visual assessment. We define our categories as follows:

0: Plumage lacking metallic luster, including plumage with uncolored (white) gloss. Examples: *Cuculus canorus, Centropus celebensis*.
1: Plumage with metallic luster, defined as a colored (of some saturation, not white) gloss. The gloss is faint-moderate in brightness/saturation. Examples: *Crotophaga major, Chrysococcyx basalis*.
2: Plumage with metallic luster that is high in brightness/saturation. Examples: *Chrycococcyx cupreus, Chrysococcyx meyeri*.

We scored the male of each species for 10 separate patches: crown, nape, mantle, rump, dorsal tail, wing covert, wing primaries/secondaries, throat, breast, and belly. We assessed the male because in cases where sexes vary in plumage coloration, the female is typically the sex lacking metallic luster. We then mapped the maximum score and mantle score for each species onto a phylogenetic tree of Cuculidae. The tree was generated by downloading 1000 trees from birdtree.org, which is based on the Jetz et al. (2012) phylogeny. We only included species that had genetic data, amounting to 126 species in total (88% of the total). From our distribution of 1000 trees, we generated a maximum clade credibility tree using TreeAnnotator (Drummond et al., 2012). Branch lengths were summarized by taking the mean branch lengths from across the distribution of time trees.

### 4.2. Feather sampling and microscopy

We sampled feathers from museum bird skin specimens (in total 21 specimens: 13 from The Academy of Natural Sciences of Drexel University and 8 from The American Museum of Natural History, details in the Appendix, §B). For each specimen, a single feather was plucked from the mantle using forceps. One species, *Phaenicophaeus diardi*, was instead sampled from the belly patch using the same technique. In addition, the sample from *Chrysococcyx xanthorhynchus* came from an unknown patch with metallic luster, since we opportunistically sampled a feather that fell off the specimen during handling. To visualize the microstructure of barbules and barbs, we imaged the dorsal surface of each feather sample using a confocal microscope (Keyence VK-X3050 Confocal).

The embedding protocol to prepare samples for transmission electron microscopy (TEM) imaging was based on methods described in Shawkey et al. (2003) with modifications based on discussions with Nicholas M. Justyn (Swansea University) and Paul Shao (Princeton University).

We cut a few barbs from each feather using a razor blade and placed the barbs in eppendorf tubes. We washed the samples by adding 100% ethanol and leaving them on a bench-top shaker for 20 minutes. This procedure was repeated once. We then mixed the embedding resin following the instructions on our kit (EMbed 812 kit, Electron Microscopy Sciences, Fort Washington, PA, USA) for a hard resin (20ml EMBed-812, 9ml DDSA, 12ml NMA, 0.72ml DMP-30). We infiltrated the feather samples with resin over four days, using solutions of the following proportions: day 1: 85% acetone and 15% resin, day 2: 50% acetone and 50% resin, day 3: 30% acetone and 70% resin, day 4: 100% resin. For each step, samples where covered with the resin solution and left for 24h at room temperature. On day five, samples where transferred to a flat embedding mould filled with resin, and the samples were then cured at 60° for 16-20 hours. The samples were first trimmed by hand with a razor blade and then with a microtome trimming knife (DiATOME trim 45°, DiATOME, Nidau, Switzerland) to prepare the block for sectioning. We sectioned the block into 72nm ultrathin sections using a Leica Ultracut UCT Ultramicrotome equipped with a diamond knife (DiATOME ultra Diamond Knife, 3mm, 45°, DiATOME, Nidau, Switzerland). Samples were transferred to 200 mesh copper grids (covered with Formvar and carbon support film) using a perfect loop (Electron Microscopy Sciences, Fort Washington, PA, USA). We stained each sample first with with UranyLess (Electron Microscopy Sciences, Fort Washington, PA, USA) and then with lead citrate (Airless bottle, Electron Microscopy Sciences, Fort Washington, PA, USA) staining solutions. A droplet of the stain was placed on Parafilm and the grid placed on top of the droplet for one minute. For the lead citrate stain, the parfilm with a droplet of stain was put in a petri dish and surrounded with NaOH pellets, to limit CO_2_ reactions. Each grid was then washed by dipping the grid 20 times in each of 3 beakers filled with CO_2_-free water, and left to dry on filter paper.

Finally, the grids were imaged using a Talos F200X Transmission Electron Microscope at an operating voltage of 200kV.

### 4.3. Measuring melanosome diameter in TEM images

We measured the diameter of melanosomes (solid rods) in TEM images using a semi-automated image analysis approach based on the MATLAB function “imfindcircles”. This would allow us to classify them as either thin solid rods (*<*190nm in diameter) or thick solid rods (*≥*190nm in diameter, Nordén et al., 2021). In detail, we first measured a large and a small melanosome in the image to set the search range for the function. We then used morphological image closing to smooth out irregularities and holes in the image, while preserving the shape and size of objects (melanosomes). The image was then transformed to a binary image using a manually adjusted threshold. Finally, circles in this processed image was detected using the “imfindcircles” function, from which diameters could be extracted. We plotted detected circles onto the original image and checked for errors by visual inspection. We also validated our method by comparing it with the results of manually measuring melanosomes using ImageJ (Abràmoff et al., 2004) for 3 random images (Appendix §C).

The melanosome diameters we recorded in Cuculidae were compared with melanosome diameters measured from samples of various plumage colors across birds. This data was compiled from previously published studies (Li et al., 2010, 2012, Nordén et al., 2019, Babarović et al., 2019). We included only samples with solid rods. Melanosome diameters were compared using a one-way ANOVA between the following categories: Cuculidae (all solid rods, both plumage with and without metallic luster), gray, black, brown, metallic luster and structural barbule. Note that metallic luster would have been described as “iridescent” in Nordén et al. (2019) and Li et al. (2012), and structural barbules as “non-iridescent” in Babarović et al. (2019).

### 4.4. Quantifying metallic luster

All species sampled for microscopy plus an additional five species where photographed using cross-polarization photography (see Appendix B for specimen and sample information). This technique allows specular and diffuse reflections from a sample to be separated using polarization filters in front of camera and light source. Since plumage with metallic luster is uniquely characterized by a strong specular reflection that is colored, we used relative specular reflection and specular saturation as a proxy for the presence of metallic luster (see Nordén et al., 2023). We followed the methods outlined in Nordén et al. (2023) to capture and process polarization images. Briefly, our set-up consisted of a Nikon D7000 camera equipped with a rotatable linear polarization filter and two light sources covered by linear polarization film. We captured images of the mantle patch of each specimen in a cross-polarized and plane-polarized configuration, with two repeats, including a Calibrite Nano ColorChecker Classic Chart in each image. Images were captured in RAW format and color calibrated using custom MATLAB code based on code and methods in Akkaynak et al. (2014) and Sumner (2014). They were then analyzed in MATLAB to calculate the relative specular reflection (specular reflection/total reflection) and the saturation of the specular reflection. The saturation was calculated as the distance to the center in a trigonal color space (RGB).

## Acknowledgements

We would like to thank The Academy of Natural Sciences of Drexel and The American Museum of Natural History for allowing us to visit and sample their ornithological collections. We are particularly grateful to Nathan H. Rice and Jason Weckstein at The Academy of Natural Sciences of Drexel, and Paul Sweet and Augie Kramer at The American Museum of Natural History for their help and advice during our visit. Many thanks to Nicholas Justyn (Swansea University), who gave advice regarding feather sample embedding and preparation, and generously shared his embedding protocol with us. John J. Schreiber and Paul Shao at the Imaging and Analsysis Center (Princeton University) trained KKN in preparing and imaging feather samples using the microtome and the TEM, and gave invaluable advice and assistance during the microscopy data collection. We would also like to thank Rosalyn Price-Waldman (Princeton University), who helped with the embedding and confocal imaging of feathers, as well as provided helpful discussion and feedback on the project during its development.

This research was made possible by a grant from the British Ornithologists’ Union to KKN, by a Packard Fellowship for Science and Engineering to MCS, and by the generosity of Eric and Wendy Schmidt by recommendation of the Schmidt Futures Polymaths program (MCS). CRC was funded by a UK Natural Environment Research Council Independent Research Fellowship (NE/T01105X/1).

## 5. Data availability statement

All data and code used in this study will be made available in a repository once published. Preceding publication data and code can be accessed at the following link: https://www.dropbox.com/sh/4lx23wnrm8no0xv/AADT-ZrBbMxR-GECvQMc-FHaa?dl=0

## 6. Author contribution statement

KKN conceived of the study and collected data, performed data analysis and wrote the first draft of the paper. All authors (KKN, CRC, FB, MCS) actively contributed to research design, interpretation of the results and editing of the manuscript during the full length of the study.

## Appendix A. Ancestral state reconstructions

We attempted to model the evolution of metallic luster in Cuculidae using Bayesian inference and continuous-time Markov models, as well as with a maximum likelihood approach. Specifically, we ran 4 different models using BayesTrait (Pagel et al., 2004, Pagel & Meade, 2006), using the mantle patch scores as input data:

(1) Markov chain Monte Carlo analysis, all rates variable (MCMC variable), i.e. all character transitions were allowed and could have different rates.
(2) Markov chain Monte Carlo analysis, restricted, where transitions had to be sequential (MCMC restricted), i.e. transitions rates 0 *→* 2 and 2 *→* 0 were set to 0.
(3) Reverse jump Markov chain Monte Carlo (RJMCMC), where the number of parameters to include in the model is determined by the analysis (Pagel & Meade, 2006).
(4) Maximum likelihood approach, all rates variable (ML variable).

In all MCMC analyses, priors were set to an exponential distribution with a mean of 10, and run for 1010000 iterations with the first 10000 iterations discarded as burn-in. These analyses generated transition rates between character states (Table A.1), which we then used to reconstruct ancestral states using simulated stochastic character maps (Bollback, 2006) implemented in the “make.simmap” fucntion in phytools (Revell, 2012).

The results of our analyses were very variable, yet many estimates had similar likelihood values (Table A.1). While the MCMC variable analysis can be discounted due to the lower likelihood values and unrealistically high transition rates, the other analyses had similar likelihood values, yet resulted in drastically different ancestral state reconstructions (Table A.1, Figure A.1). For example, two different solutions in the RJMCMC analysis with similar likelihood suggest that Cuculidae was either state “2” at the root, and this state has subsequently been lost in every single clade (transitions to “2” is 0, Table A.1, Figure A.1C)—or state “2” was independently gained in every single species of *Chrysococcyx* (transitions to “2” is unrealistically high, Table A.1, Figure A.1D). The MCMC restricted and the ML analysis give somewhat more evolutionary plausible results, but the likelihood of these analyses are very similar to that of the RJMCMC runs (note also that the higher number of free parameters in the ML analysis is expected to increase likelihood). Thus, it is not possible on the basis of these analysis to identify a single set of well-supported transition rates. Of course, we can use our knowledge of how metallic luster is likely to have evolved in Cuculidae (e.g. based on phylogenetic bracketing) and then pick the transition rates that are most consistent with our reasoning—but this approach would not test the evolution of this trait in an unbiased way (which is our goal). With the number of parameters in these models, any evolutionary scenario could be reconstructed with enough modifications.

We conclude that the evolution of metallic luster in plumage is not well modeled by a Markov process (transition rates may not be equal across the tree, see similar issue documented by Wiens et al., 2007), and/or our data set is too small to achieve robust results.

**Table A.1.** Transition rates and root posterior probabilities for characters “0” (no metallic luster), “1” (faint-moderate metallic luster) and “2” (intense metallic luster) in Cuculidae (mantle patch), found using four different models. To demonstrate that the model runs did not find a single solution, we have included two solutions (iterations in the chain) for each MCMC model, which have similar likelihood but very different transition rates. log Lh denotes log likelihood. Model abbreviations: MCMC var, Markov chain Monte Carlo model with variable rates; MCMC res, Markov chain Monte Carlo model with rates 0 *→* 2 and 2 *→* 0 set to 0; RJMCMC, reverse jump Markov chain Monte Carlo model; ML, maximum likelihood. See text for further explanation of models.

| Model | log Lh | $0 \rightarrow 1$ | $0 \rightarrow 2$ | $1 \rightarrow 0$ | $1 \rightarrow 2$ | $2 \rightarrow 0$ | $2 \rightarrow 1$ |
| --- | --- | --- | --- | --- | --- | --- | --- |
| MCMC var | -103.0 | 0.3078 | 0.1202 | 0.3781 | 1.851 | 2.771 | 21.06 |
|  | -103.1 | 11.04 | 1.878 | 15.20 | 1.714 | 10.63 | 32.83 |
| MCMC res | -84.76 | 0.02396 | 0 | 0.03461 | 0.007663 | 0 | 0.1014 |
|  | -87.74 | 0.02416 | 0 | 0.03082 | 0.6918 | 0 | 7.307 |
| RJMCMC | -85.90 | 0.02806 | 0 | 0.02806 | 0 | 0.02806 | 0.02806 |
|  | -86.13 | 0.01667 | 0 | 0.01667 | 0.01667 | 0.01667 | 0.01667 |
| ML | -84.15 | 0.02147 | 0.001531 | 0.03199 | 0.001347 | 0.000 | 0.06087 |

| Model | $p$ (root 0) | $p$ (root 1) | $p$ (root 2) |
| --- | --- | --- | --- |
| MCMC var | 0.3333 | 0.3333 | 0.3333 |
|  | 0.3333 | 0.3333 | 0.3333 |
| MCMC res | 0.3017 | 0.3449 | 0.3534 |
|  | 0.3489 | 0.3256 | 0.3255 |
| RJMCMC | 0.000105 | 0.000084 | 0.9998 |
|  | 0.7823 | 0.1397 | 0.07799 |
| ML | 0.3544 | 0.3770 | 0.2686 |

**Figure A.1.**
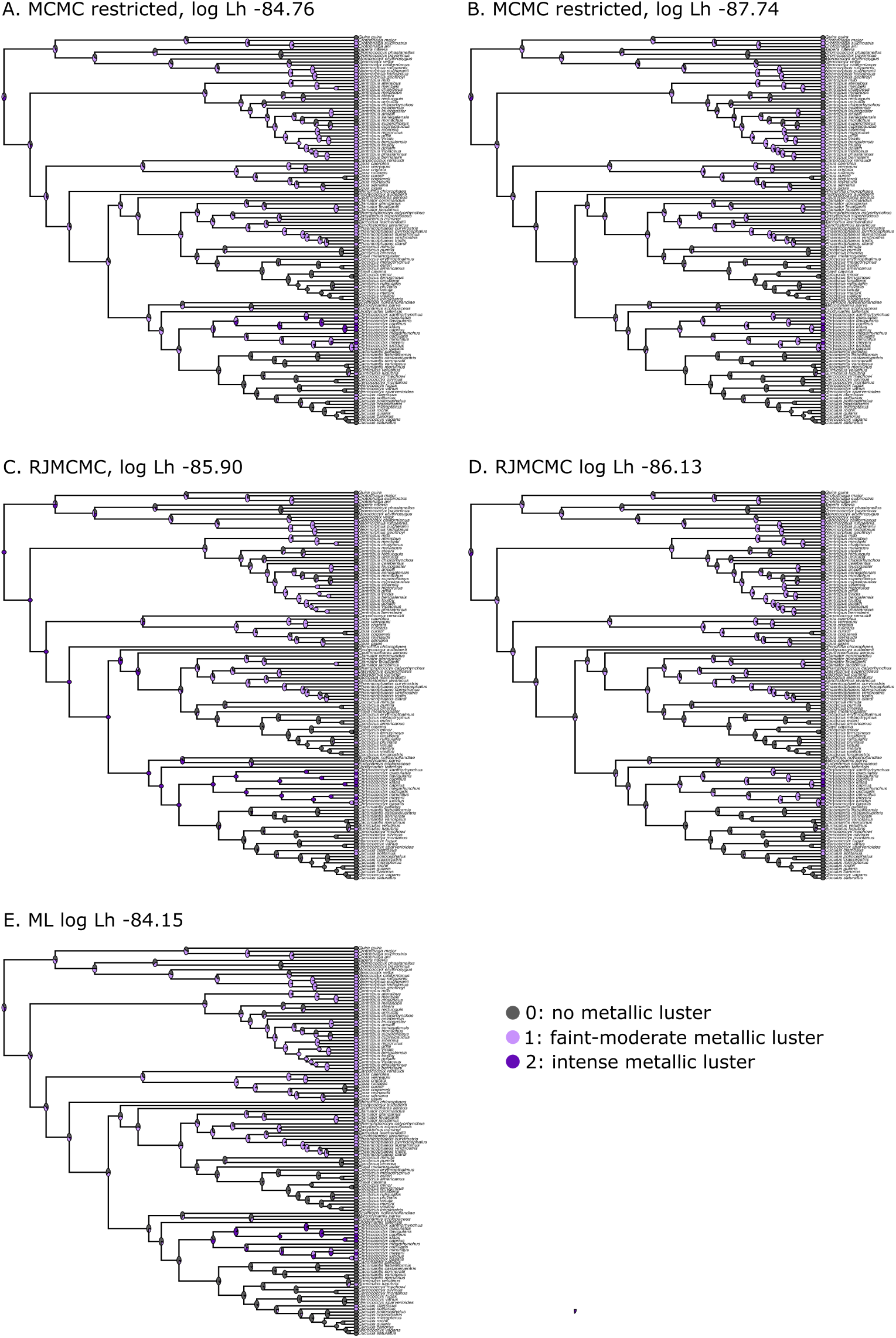
Ancestral state reconstructions of metallic luster in Cuculidae (mantle patch) using different transition rates extracted from the models (Table A.1). The pie charts at the nodes represent the percentage of iterations that a node was reconstructed as a particular character state in the stochastic character mapping. Note that the RJMCMC model presents two extreme but quite unlikely scenarios, which nevertheless have a similar likelihood values to the ML and MCMC restricted analysis.

## Appendix B. Specimens sampled and imaged with cross-polarization photography

**Table B.1.**
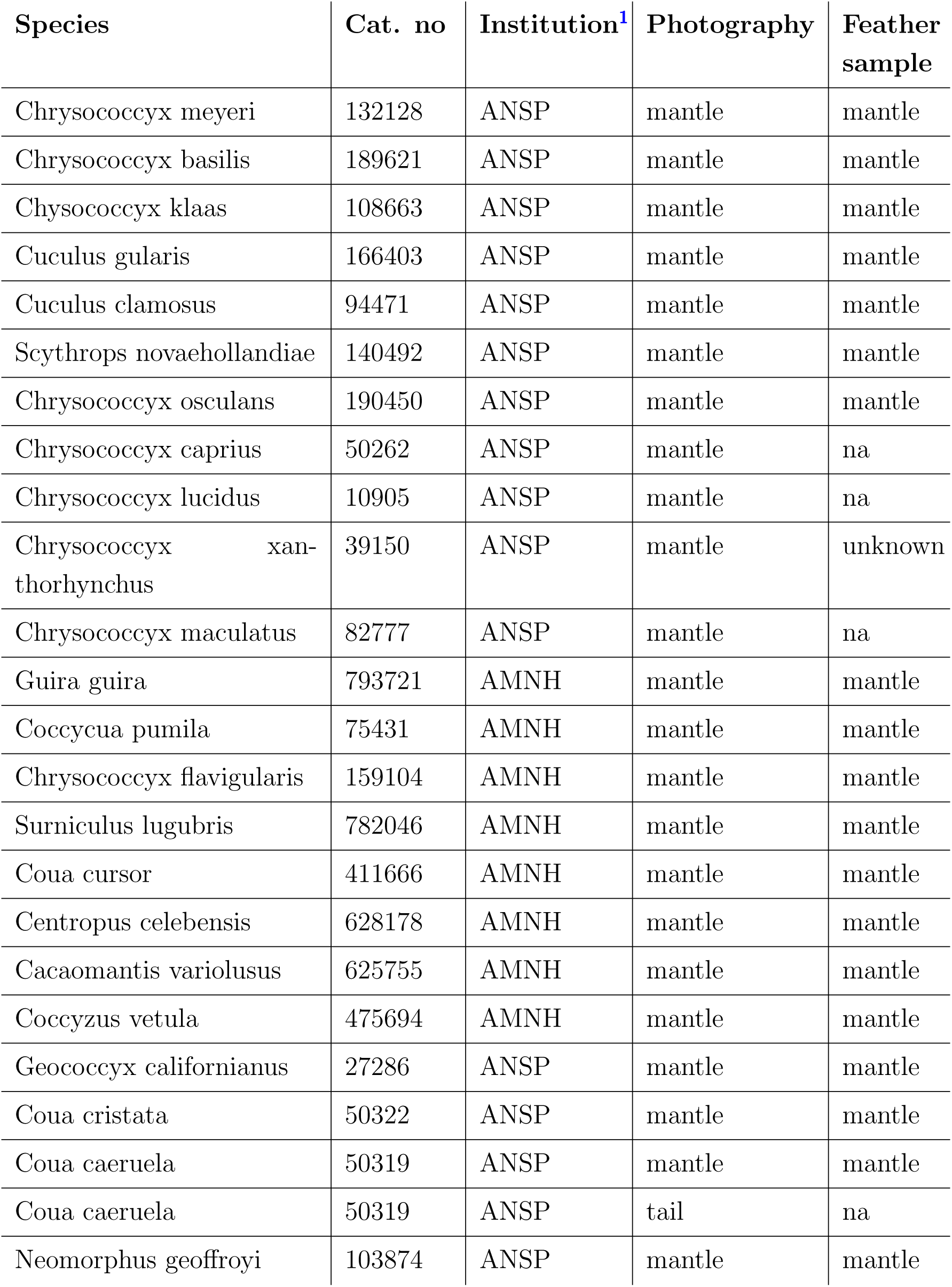
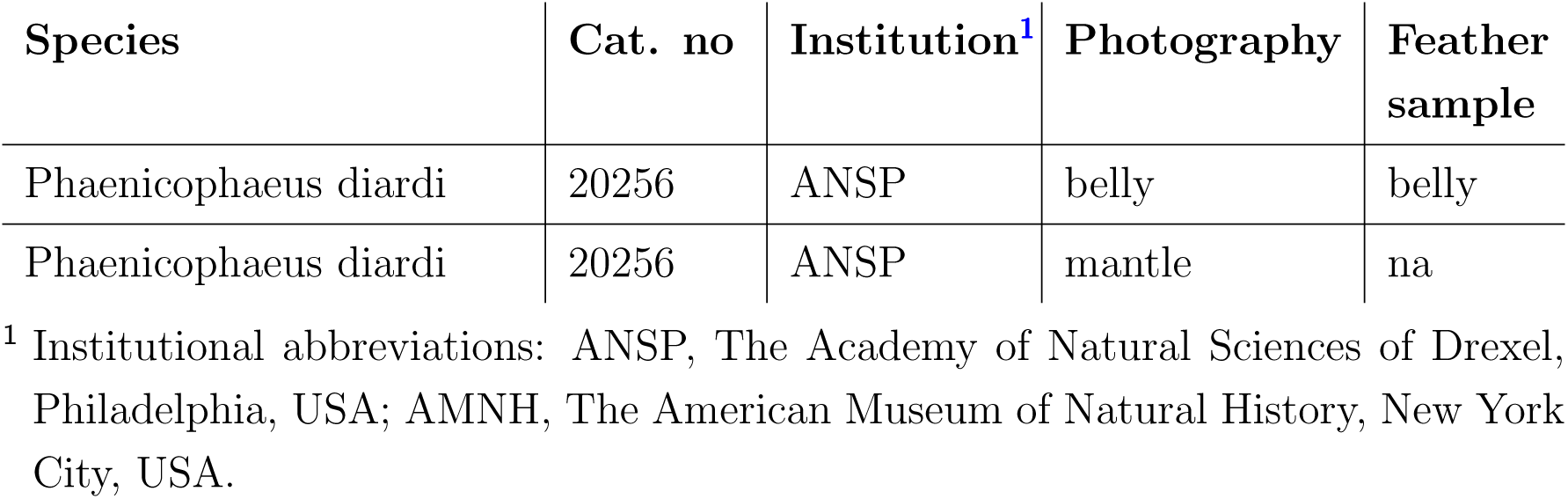
Specimens sampled and/or photographed for the study.

## Appendix C. Validation of image analysis method to estimate melanosome diameters

**Figure C.1.**
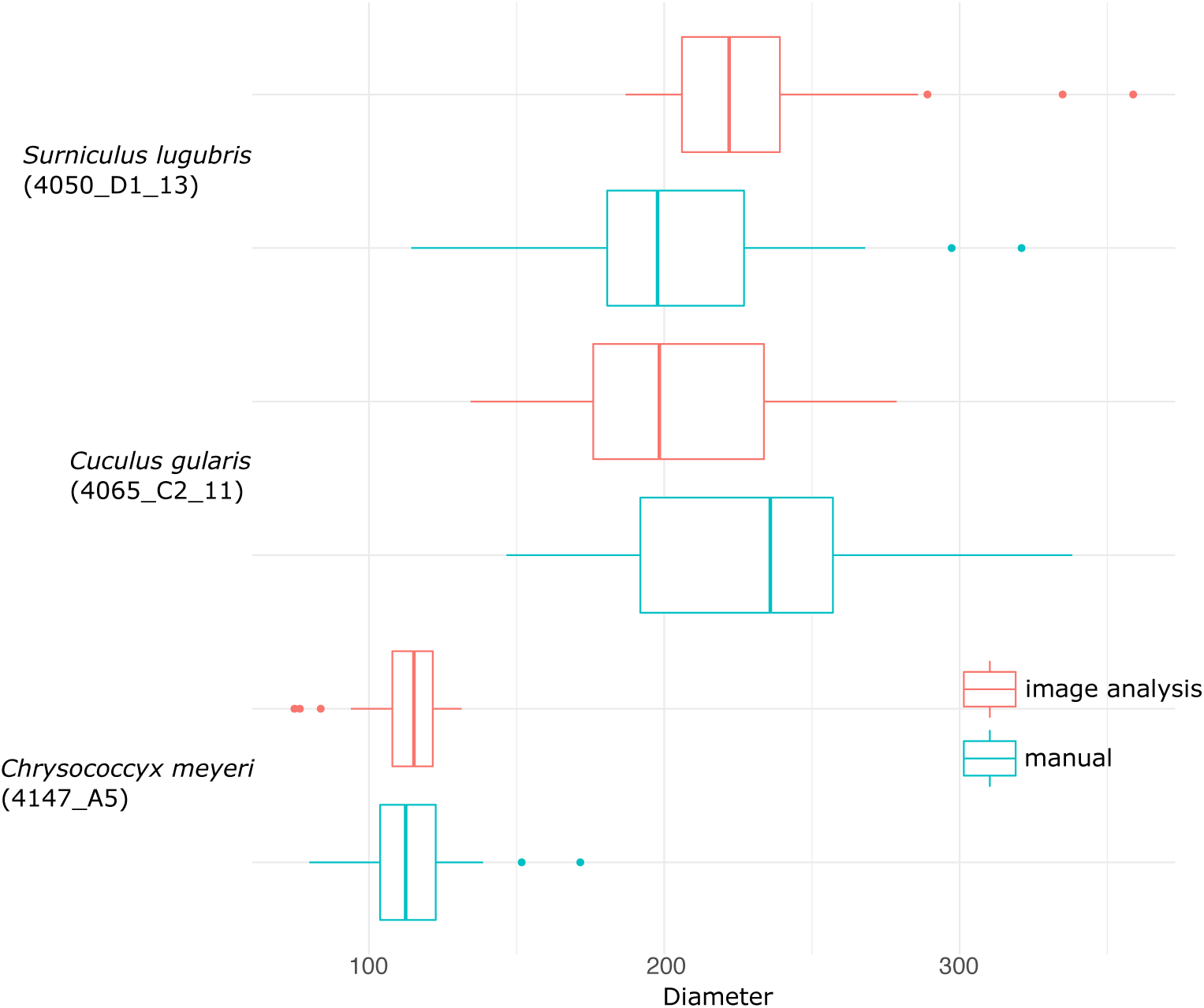
Comparison of melanosome diameter measurements using the semi-automated image analysis method (red) and manual measuring by hand using ImageJ (blue), for three TEM images (each from a different species). The code under species names identifies the specific sample and image measured.

